# New distribution data and phylogenetic approach reveal bioregionalization of the western Palearctic ants

**DOI:** 10.1101/2022.03.10.483749

**Authors:** Runxi Wang, Jamie M. Kass, Christophe Galkowski, Federico Garcia, Matthew T. Hamer, Alexander Radchenko, Sebastian Salata, Enrico Schifani, Zalimkhan M. Yusupov, Evan P. Economo, Benoit Guénard

**Affiliations:** School of Biological Sciences, The University of Hong Kong, Kadoorie Biological Sciences Building, Pok Fu Lam Road, Hong Kong SAR, China; Biodiversity and Biocomplexity Unit, Okinawa Institution of Science and Technology Graduate University, Onna, Okinawa, Japan; Société Linnéenne de Bordeaux, 1 place Bardineau, 33000, Bordeaux, France; Iberian Myrmecological Association, Barcelona, Spain; Schmalhausen Institute of Zoology of the National Academy of Sciences of Ukraine, 15 Bogdana Khmelnitskogo str., 01030, Kiev, Ukraine; Department of Biodiversity and Evolutionary Taxonomy, University of Wroclaw, Przybyszewskiego 65, PL-51148 Wroclaw, Poland; Department of Chemistry, Life Sciences and Environmental Sustainability, University of Parma, Parco Area delle Scienze 11/a, 43124 Parma, Italy; Tembotov Institute of Ecology of Mountain Territories of the Russian Academy of Science, I. Armand Street 37-A, Nalchik city, 360051, Kabardino-Balkarian Republic, Russia; Radcliffe Institute of Advanced Study, Harvard University, Cambridge, MA, USA

**Keywords:** biogeography, regionalization, beta diversity, Europe, Formicidae, insect, species distribution modelling, beta diversity, phylogenetic turnover

## Abstract

**Aim:** Biogeographic regionalization has fascinated biogeographers and ecologists for centuries and is endued with new vitality by evolutionary perspectives. However, progress is scant for most insect groups due to shortfalls in distribution and phylogenetic information, namely Wallacean and Darwinian shortfalls respectively. Here, we used the western Palearctic ants as the case to tackle these shortfalls and test their biogeographic structure through novel distribution data and phylogenetic approaches.

**Location:** Western Palearctic realm.

**Taxon:** Ants (Formicidae).

**Methods:** Firstly, we developed a refined database integrating the occurrences of 747 ant species across 207 regions of the western Palearctic realm, based on newly expert-validated records derived from the existing global ant biodiversity informatics. Using range estimates for these species derived from polygons and species distribution modelling, we produced species assemblages in 50 × 50 km grid cells. We calculated taxonomic and phylogenetic turnover of ant assemblages, performing hierarchical clustering analysis using the Simpson dissimilarity index to delineate biogeographic structure.

**Results:** At both the regional list- and grid assemblage-levels, the Mediterranean has higher turnover and more biogeographic regions than northern Europe, both taxonomically and phylogenetically. Delineations based on grid assemblages detected more detailed biogeographic transitions, while those based on regional lists showed stronger insularity in biogeographic structure. The phylogenetic regionalization suggested closer but varied affinities between assemblages in comparison to the taxonomic approach.

**Main conclusions:** Here, we integrated expert-validated regional lists, species distribution modelling, and a recent phylogeny to tackle Wallacean and Darwinian shortfalls for an important insect group by developing a next-generation map of biogeographic regionalization for the western Palearctic ants. The results of this study suggest strong constraints from geographic barriers and potential effects of climatic history on ant distributions and evolutionary history, and also provide baseline spatial information for future investigations of regional insect distributions.

## INTRODUCTION

Organisms are not uniformly distributed on Earth—most are restricted to particular areas due to ecological and evolutionary processes, ultimately forming distinct biogeographic regions (Wallace, 1876; Lomolino et al., 2016). The concept of bioregionalization is thus fundamental to the classification of species distributions (Wallace, 1894; Kreft & Jetz, 2010; Morrone, 2018) and provides a powerful framework for informing empirical studies in ecology and evolution, as well as applied studies in conservation and management (Wallace, 1876; Kreft & Jetz, 2010; Holt et al., 2013). For centuries, biogeographers have proposed various regionalization systems mainly based on plants and terrestrial vertebrates (e.g., Wallace, 1876; Smith, 1983; Cox, 2001; Kreft & Jetz, 2010; Holt et al., 2013), but similar classification efforts based on insects, which represent an overwhelming component of biodiversity (Stork, 2018), have been scant.

This taxonomic bias in regionalization may raise potential issues for the general representativity of described biogeographic patterns. First, insects have evolved complex and unique life histories and strategies (e.g., parasitism) (Gullan & Cranston, 2014). Moreover, mounting evidence has shown that since the late Quaternary, the distributions of many vertebrate groups have been shaped substantially by targeted anthropogenic selection pressures like hunting and domestication (Faurby & Svenning, 2015; Santos et al., 2020), which for the most part have not historically had direct effects on insects. Instead, likely direct drivers of insect distributions over this time period have been climate, geology, and land cover change (Elias, 1991; Rueda, Rodríguez & Hawkins, 2010; Ballesteros-Mejia et al., 2017). Due to these differences, expanding our knowledge of biogeographic patterns to include insects would enhance, and potentially validate, our current understanding of biogeographical regionalization.

The lack of detailed distributional information for insects (i.e., Wallacean Shortfall, Lomolino, 2004; Guénard et al., 2017), however, represents a major challenge to studying insect regionalization. Most of the existing insect regionalization systems are based on regional lists, which represent the most widely used geographic units and the most comprehensive distributional information for insect taxa (e.g., Dennis, Williams & Shreeve, 1991; Heiser & Schmitt, 2010; Vitali & Schmitt, 2017). These regional lists provide useful biogeographical information, but their geographic units are normally taxon-specific, delineated as expert range polygons with varying detail and accuracy that are limited to sampled areas. Therefore, their boundaries and ranges may lack quantitative validation and be limited by sampling bias, resulting in regionalization patterns with potential limitations for cross-taxa comparisons (Rueda et al., 2010). Species distribution models (SDMs), which estimate relationships between species’ occurrence localities and environmental variables to make predictions of range extents, provide a potential solution for tackling the Wallacean Shortfall for taxa with few distributional data such as insects (Diniz-Filho, De Marco Jr, & Hawkins 2010; Ballesteros-Mejia et al., 2017, Kass et al. 2020). Species distribution modeling represents a data-driven approach to generating reproducible range estimates and making predictions for unsampled areas (Peterson et al. 2018). Although SDMs for low-data species in particular can be susceptible to sampling bias, statistical overfitting, and other methodological issues (Galante et al., 2017), recently proposed methods for tuning model complexity (Radosavljevic & Anderson 2014) and accounting for sampling bias (Phillips et al. 2009) can help remedy many of these issues.

Along with a lack of distributional information for insects, there is also a dearth of phylogenetic information (i.e., the Darwinian Shortfall, Lomolino, 2004), representing another big challenge for understanding insect biogeography. Historically, the delineation of biogeographic regions has been based solely on taxonomic information, but in recent years, progress in phylogenetic methods (i.e., phylogenetic regionalization) has provided new opportunities to explore evolutionary relationships for entire assemblages and enhance the objectivity and repeatability of delineations (Holt et al., 2013; Daru et al., 2017; Ye et al., 2019). Especially for taxonomically challenging and complex groups like insects, the phylogenetic approach is attractive because it provides an inclusive measurement for taxonomic classification based on phylogenetic distance. However, the role that evolutionary history has played in regionalization remains rarely explored for insects (Diniz-Filho et al., 2010).

Ants (Formicidae) represent good model organisms to investigate the biogeography of insects due to their wide distributions across climates and biomes, but also because of their varied and dominant roles in ecosystems as keystone species (Hölldobler & Wilson, 1990; Guénard et al., 2017). Modern biodiversity informatics and recent research on ant macroevolution have resulted in new data on their distributions and phylogeny (Economo et al., 2018; Guénard et al., 2017; Kass et al., submitted). Although new species are continuously being described across the globe and significant effort directed towards sampling and identification remains needed, ants within the West Palearctic realm are relatively well-documented and are arguably among the best-known ant faunas on the globe.

The biogeography of the western Palearctic realm has been studied for centuries and its insect distributions are thought to have been shaped by climate change since the Last Glacial Maximum. Climate refugia in the Mediterranean region harbor high species endemism and show high heterogeneity in species composition, while species distributions in the more northern regions have a more homogenous structure (Dennis et al., 1991; Fattorini & Ulrich, 2012; Vitali & Schmitt, 2017) and are suggested to be driven by postglacial dispersal processes (Hewitt, 1999; Schmitt, 2007; Calatayud et al., 2019). Previous western Palearctic regionalization systems have mainly focused on a few taxa including butterflies (e.g., Dennis et al., 1991; Rueda, Rodrgí uez & Hawkins, 2010), dragonflies (e.g., Heiser & Schmitt, 2010; Heiser, Dapporto & Schmitt, 2014) and some beetle taxa (e.g., Fattorini & Ulrich, 2012; Vitali & Schmitt, 2017). All of them, however, lack fine-scale distributional data or a phylogenetic understanding. Here, we developed a novel dataset including fine-scale regional lists, SDMs, and a large-scale phylogeny of the western Palearctic ants to tackle these shortfalls and to delineate the most comprehensive regionalization of insects in the western Palearctic realm to date (Figure S2.1).

First, we evaluated the biogeographic structure of western Palearctic ants based on both regional lists and grid assemblages generated from SDMs. We expected that both sets of results would have similar spatial patterns, but that the grid assemblage delineation would better identify transitions between biogeographic regions and the uniqueness of ant assemblages. We also expected the SDM results would help fill in gaps by making predictions for areas with limited sampling, thus reflecting more objective spatial extents of ant distributions. Secondly, we tested whether the phylogenetic regionalization results in different patterns from those obtained with the taxonomic approach. A historical connection between regions based on shared evolutionary history may fail to be detected under a taxonomic delineation (Ye et al., 2019). For example, allopatric speciation events driven by geographical isolation can increase the dissimilarity of species composition between two regions yet a phylogenetic affinity between them would remain (Daru et al., 2017). We thus expect the relationships between regions resulting from phylogenetic regionalization to differ from those of the taxonomic delineation, especially for geographically isolated species, and that these differences will help to reveal a more multilayered regional evolutionary history for the western Palearctic ants.

## METHODS

### Ant distributions and phylogeny

#### European Ant Distribution (EUAD) database

We developed the EUropean Ant Distribution (EUAD) database, a new collection of ant species’ occurrence data for the western Palearctic realm. This database is derived from the Global Ant Biodiversity Informatics (GABI, Guénard et al., 2017) but features a higher spatial resolution for the region. Ultimately, we compiled native ant taxa occurrence information for each of the 207 geographic divisions (i.e., regional lists) for the western Palearctic realm, which was previously divided to 57 regions in GABI. Our definition of the western Palearctic realm does not include North Africa and the Arabian Peninsula because of historically poor sampling and the lack of recent taxonomic revisions for species in those regions. Geographical divisions used in the database were delimited based on either administrative region (GADM, version 2.8, accessed 1st Sep. 2020) or modified areas based on the physical geographic area (e.g., islands and mountains), depending on data availability (details can be found in Figure S2.2).

#### Validation of regional lists

Preliminary versions of the database were validated by ant experts (co-authors of this study) who identified dubious records and provided additional information (e.g., unpublished or missing records) to complete and provide more accurate ant range maps. Occurrence records were deemed dubious for reasons including nomenclatural changes in recent taxonomic revisions, outdated taxonomy, and misidentifications (which can be numerous in older literature or databases). Ultimately, the validation process showed that 8% of species incidence records in the preliminary versions were dubious (and either corrected or excluded for later analysis) and also contributed 16% new incidence records to the final database (Figure S2.3). For all ant taxa in our database, we also verified nomenclature based on AntCat, an online, global catalog of ants (Bolton, 2021), with validation and inclusion of taxa up to July 1^st^ 2021. Here, we treated valid subspecies as species in our analysis, which resulted in a total of 747 valid native species (including 40 subspecies) for regional lists (see Table S1.1).

#### Grid assemblages

We first made grid-based estimates of the western Palearctic ant assemblages using range estimates from SDMs developed for a global analysis on ant diversity (Kass et al., submitted). All analyses were performed using the statistical computing language R 4.0.2 (R Core Team, 2020). Ranges were estimated for low-data species (<5 occurrence records) with univalue polygons (either buffered [30 km] points or convex/alpha hull, depending on data availability), and for species with sufficient data (≥ 5 occurrence records) using SDMs. We used the presence-background machine-learning algorithm Maxent to train models over a study extent defined by their polygon range estimate (buffered alpha hull) using 19 bioclimatic predictor variables at 10 arcminute resolution (∼20 km at the equator) from Worldclim 2.0 (Fick & Hijmans 2017). We tuned models for optimal complexity (i.e., combinations of feature classes and regularization multipliers) using sequential criteria of cross-validation results (based on the 10 percentile omission rate and validation AUC; Radosavljevic & Anderson 2014) with the R package *ENMeval* 2.0.0 (Kass et al. 2021). We used these tuned models to make predictions of suitability over the species’ study extents, effectively constraining range estimates to the limits of the occurrence data, and made them binary (presence/absence predictions) by thresholding with the 10 percentile omission value. Range estimates represented by polygons for low-data species were converted to 10 arcminute grid cells to align with the modeled range estimates. We then projected the range estimates for all species to a 50 × 50 km resolution equal -area extent (Albers Equal Area Conic Projection) covering our study area in the wetern Palearctic realm. Binary model predictions were projected to this coarser resolution using bilinear interpolation that selected the maximum neighborhood value, resulting in predicted presence for 50 km × 50 km cells when any underlying 10 arcminute (∼20 km) cells had predictions of presence. Grid-based analyses used the R packages *raster* (Hijmans, 2021) and *sf* (Pebesma, 2018). We excluded grid cells with fewer than 5 species to control for the negative influence of low species richness in the analysis (He et al., 2020). Ultimately, we obtained 4527 grid assemblages representing 711 species (36 species were not included due to insufficient occurrence data in our study area, Table S1.1 and Figure S2.4).

#### Phylogeny

We derived phylogenetic information from a recently reconstructed, large-scale ant phylogeny that represents most of currently recognized genera and provides valid relationships between genera and their associated uncertainties (Economo et al., 2018). We updated the nomenclature of taxa in the 100 posterior phylogenetic trees and pruned those trees based on the list of native taxa in the western Palearctic realm. This process used the R packages *geiger* (Slater et al., 2012) and *picante* (Kembel et al., 2010). As a result, estimated phylogenetic information for 641 species (86% of the total) was available for further analysis (see Figure S2.4).

### Measurement of turnover

We determined the uniqueness of ant assemblages by measuring turnover of taxonomic and phylogenetic compositions using the Simpson’s index dissimilarity metric on assemblage pairs (Baselga, 2010; Kreft & Jetz, 2010). We calculated two pairwise distance metrics: taxonomic dissimilarity (βsim) and phylogenetic dissimilarity (Pβsim) with the following formula:

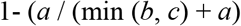

For βsim, *a* is the total number of species shared between two assemblages, and *b* and *c* are the numbers of species unique to each assemblage. For Pβsim, *a* is the total branch length shared between assemblages, and *b* and *c* are the lengths of unique branches to each assemblage (Daru et al., 2017; Holt et al., 2013). Pβsim was based on the median value across 100 posterior trees to control the uncertainty of phylogeny. Higher or lower values of βsim or Pβsim indicate the assemblages are less or more similar to each other, respectively. The mean pairwise βsim or Pβsim of each assemblage was calculated to show spatial patterns, where higher βsim or Pβsim denotes higher uniqueness of species composition? or evolutionary history for an assemblage. We also used null models to test if the observed assemblages were more or less similar (taxonomically and phylogenetically) than expected by chance. The standardized effect size (SES) of turnover was calculated as:

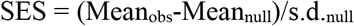

where Mean_obs_ is the mean of the observed βsim or Pβsim, and Mean_null_ and s.d._null_ are the mean and standard deviation of the null distribution for each randomized assemblage. The null model used the Independent Swap algorithm to control for the effects of both the species pool and different richness of assemblages (Gotelli, 2000) and was run for 1000 iterations. Higher or lower values of SES here indicate that the assemblages have higher or lower turnover than expected, respectively. The calculations for turnover and the null model used the R package *betapart* (Baselga et al., 2021). The difference of pairwise βsim and Pβsim distance matrices were assessed using the Mantel correlation test in the R package *vegan* (Oksanen et al., 2020). The spatial congruence between the average βsim and Pβsim of assemblages was measured using a modified t-test that can control for spatial autocorrelation by correcting the degree of freedom (Clifford et al., 1989), using the R package *SpatialPack* (Vallejos, Osorio & Bevilacqua, 2020).

### Delineation of biogeographic structure

We performed clustering analysis based on pairwise taxonomic and phylogenetic turnover for the biogeographic structure classification. To choose the most appropriate clustering algorithm for our data, we examined eight candidates: 1) unweighted pair-group method using arithmetic averages (UPGMA); 2) unweighted pair-group method using centroids (UPGMC); 3) Ward’s method (WARD); 4) single lineage (SL); 5) complete lineage (CL); 6) weighted pair-group method using arithmetic averages (WPGMA); 7) weighted pair-group method using centroids (WPGMC); 8) divisive hierarchical clustering (DIANA). We measured algorithm performance by calculating the cophenetic Pearson correlation and the Gower distance to test the degree of data distortion in models (Gower, 1983; Holt et al., 2013; Legendre & Legendre, 2012), leading us to select UPGMA for our analysis (Table S2.2).

Based on the dendrograms from the UPGMA analysis, we used three different metrics to choose the optimal classification of biogeographic regions and subregions: 1) average silhouette width (ASW) with the R package *cluster* (Maechler et al., 2021); 2) Kelly-Gardner-Sutcliffe penalty (KGS) with the R package *maptree* (White & Gramacy, 2012); and 3) Bootstrap mean instability (Kelley et al., 1996; Kreft & Jetz, 2010; Legendre & Legendre, 2012) with the R package *fpc* (Hennig, 2020). High reliability of clusters is indicated by higher ASW and lower KGS, and mean instability. We defined the biogeographic regions as regions characterized by distinct and coherent ant assemblages that can be delineated clearly in space (Kreft & Jetz, 2010; Morrone, 2018; He et al., 2021). We also used the R package *phyloregion* (Daru, Karunarathne & Schliep, 2020) to visualize regionalization patterns.

## RESULTS

### Spatial turnover of West Palearctic ants

Taxonomic and phylogenetic turnover (βsim and Pβsim) showed high correlation and spatial congruence in both regional lists and grid assemblages (Pearson’s correlation R = 0.87∼0.96, *P* < 0.001, Table S2.3). In regional lists, the highest turnover values for both metrics were observed in the Mediterranean peninsulas and islands, and southern Anatolia, while central (e.g., Alps and Carpathians) and northern Europe and the British islands showed lower βsim (Figures 1a and e). The observed βsim and Pβsim was higher than expected in the southern Iberian, southern Anatolia and some Mediterranean islands such as Sicily, Crete and Cyprus (Figures 1b and f). The spatial turnover of grid assemblages presented a similar pattern: both turnover metrics were highest in the south of the western Palearctic realm and decreased poleward (Figures 1c and g). The standardized effect size (SES) also suggested the observed βsim and Pβsim of grid assemblages were significantly higher than the random patterns in the southwestern Palearctic realm, similar to the results from regional lists (Figures 1d and h).

**Figure 1.**
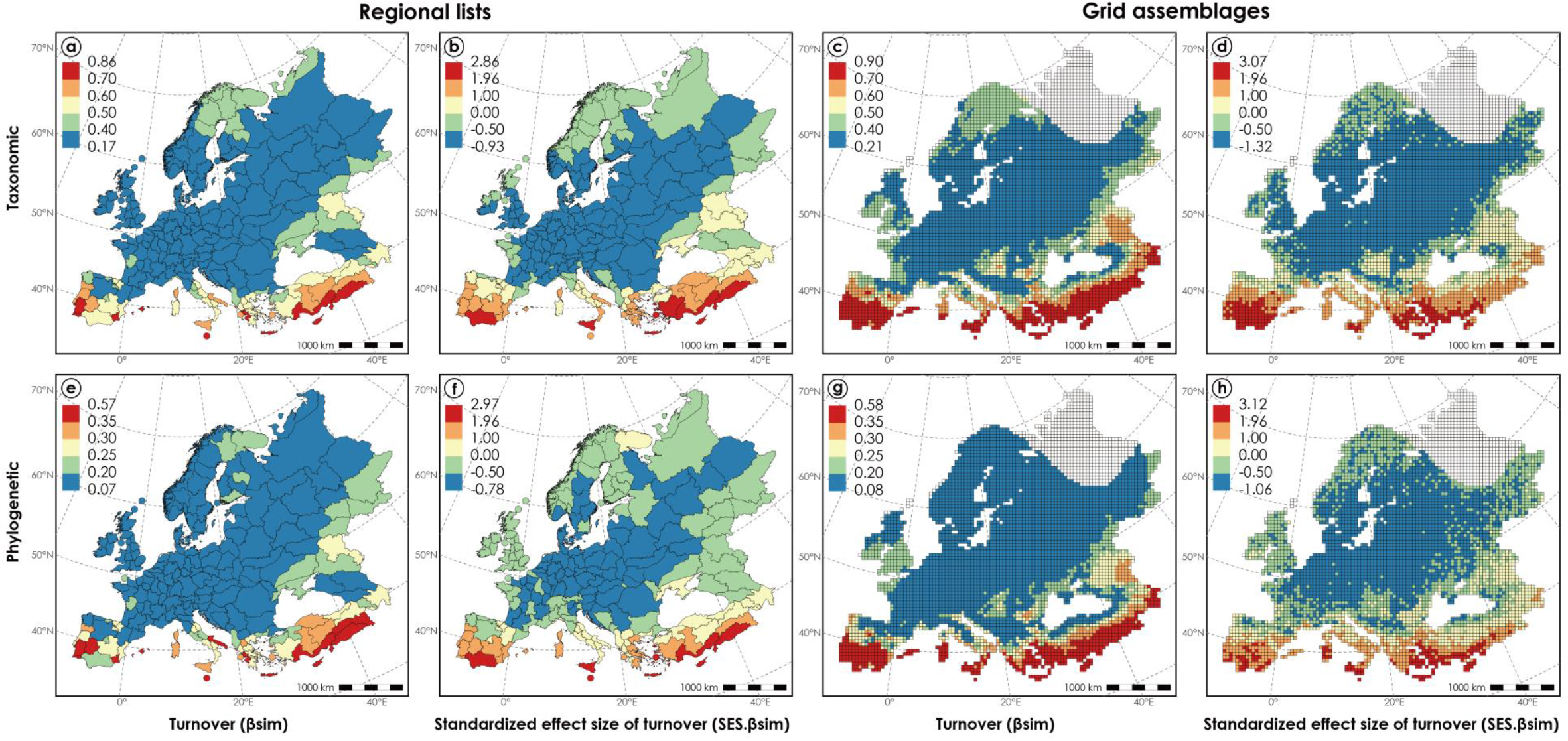
Spatial patterns of turnover of the western Palearctic ant assemblages. Taxonomic (a-d) and phylogenetic (e-h) turnover of regional lists (a, b, e, f) and grid assemblages (c, d, g, h). Both observed average value of taxonomic (βsim) (a, c) and phylogenetic (Pβsim) (e, g) turnover and their standardized effect size (SES) results from randomization test (Independent Swap) were calculated (b and d for SES.βsim; f and h for SES.Pβsim). Values that are not available were indicated by the white colour on the map.

### Biogeographic structure of West Palearctic ants

#### Delineation of regional lists

The hierarchical clustering based on pairwise βsim of regional lists suggested six biogeographic regions and four subregions (Figures 2a and b; Figure S2.5): (1) Sicily and Maltese islands (SM) were closely grouped and distinct from the rest of the western Palearctic realm; (2) Cyprus (CY), (3) Southeastern Anatolian (SEA) and (4) Eastern Mediterranean (EM) including Turkey (EM1), Aegean islands and Balkans (EM2), were grouped together; (5) Western Mediterranean (WM) included the Iberian Peninsula and the Balearic Islands; (6) European region (EU) including Apennine Peninsula, Adriatic Balkans, Corsica and Sardinia islands (EU1) and European mainland and the British Isles (EU2), were grouped in the same cluster. The western Mediterranean and European regions were grouped as the sister cluster of the Eastern Mediterranean region, Southern Anatolian and Cyprus.

**Figure 2.**
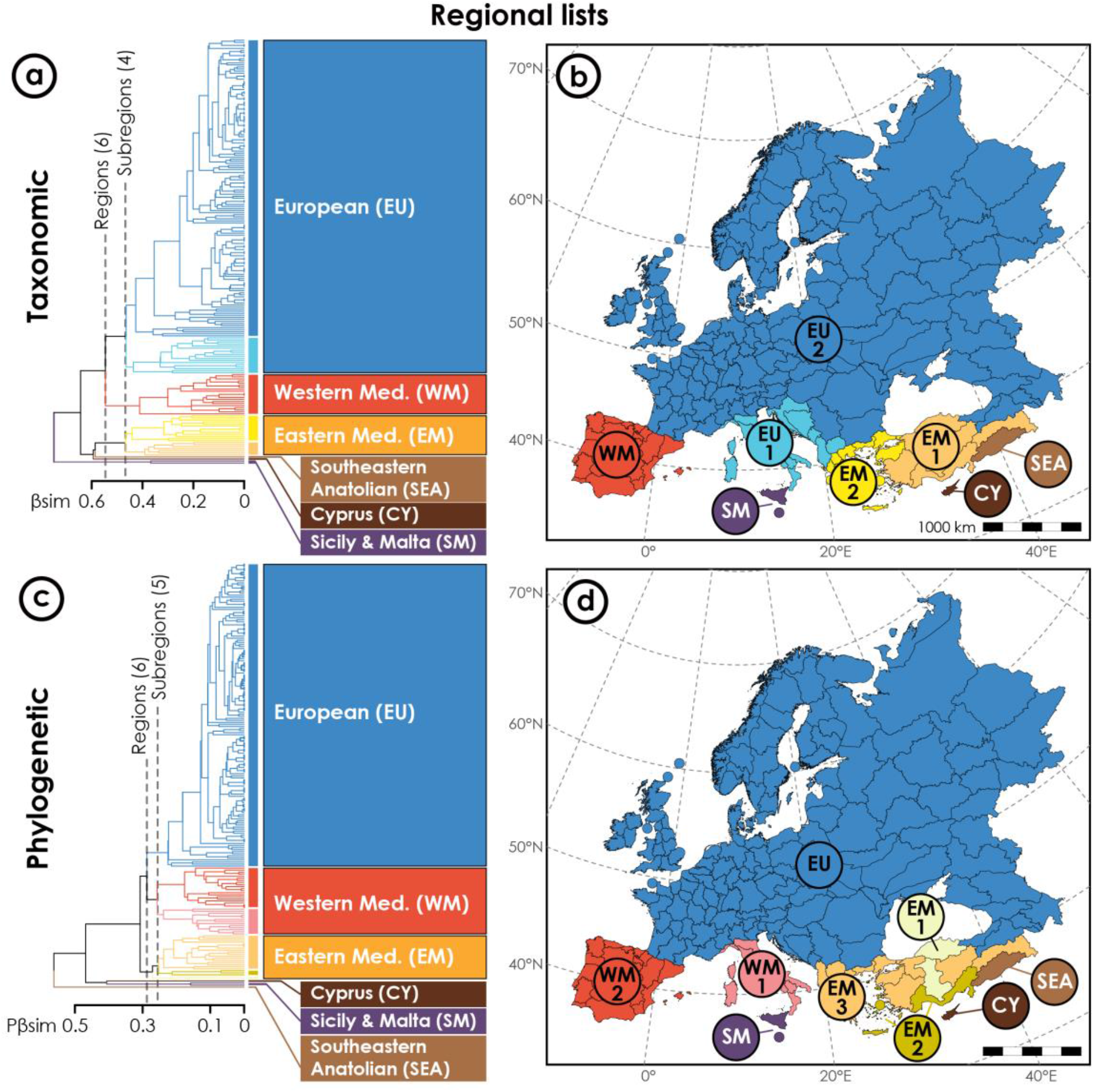
Regionalization of the western Palearctic ant fauna based on regional lists. Dendrograms (a, c) and maps (b, d) resulting from the unweighted pair-group method using arithmetic average (UPGMA) hierarchical clustering based on βsim (a-b) and Pβsim (c-d) matrices. Colours used to characterize particular regions in dendrograms and maps are identical.

The delineation based on Pβsim also recognized six biogeographic regions and presented similar spatial patterns compared to the results of βsim (Figures 2c and d; Figure S2.5). However, biogeographic affinities between some regional assemblages were different in the phylogenetic delineation. The Apennine Peninsula, Corsica and Sardinia islands (WM1) were identified as the sister group of the Iberian Peninsula and the Balearic Islands (WM2), and a part of the Western Mediterranean (WM). The western Anatolian (EM1), Aegean islands and Mediterranean Turkey (EM2) were distinct from the rest of the Eastern Mediterranean (EM). The Adriatic part of the Balkans was assigned to be a part of the European region (EU). The Southeastern Anatolia (SEA), Sicily and Maltese (SM) and Cyprus (CY) islands were suggested as the outgroup of the rest of the western Palearctic.

#### Delineation of grid assemblages

Grid assemblage clustering consistently recognized three biogeographic regions of western Palearctic ants: Western Mediterranean (WM), Eastern Mediterranean (EM) and European region (EU) (Figure 3; Figure S2.6). There were six and seven subregions detected based on βsim and Pβsim, respectively. The Western Mediterranean region included Corsica, Sardinia, Sicily and Maltese islands (WM1, Figure 3d) and also included the Mediterranean coast of France (WM2, Figure 3d) in the phylogenetic but not in the taxonomic delineation (Figure 3b). The Eastern Mediterranean region extended from the Eastern Caucasus (ECA, Figures 3b and d) to the Balkan Peninsula and even reached the Apennine Peninsula and Corsica island (EM3, Figure 3b) in the taxonomic delineation. Crete island (EM2, Figure 3b) was suggested to be distinct from the rest of the Eastern Mediterranean region based on βsim while the southern Balkans, Aegean regions, southern Anatolia and Cyprus were grouped together based on Pβsim (EM1, Figure 3d). The Western and Eastern Mediterranean regions were grouped as the sister cluster of the European region. Some lowland areas in the south of the European continent were identified as a subregion (EU1) of the European region (EU, Figure 3): lowland in the north of Caucasus, steppe in the north of Black Sea, plains in the south of Carpathian and Alps mountains and in the north of Balkan, Dinaric Alps and Apennine mountains. This subregion also included southern France and northern Iberia, except the Pyrenees, in the taxonomic delineation (Figure 3b) while the Apennine Peninsula and Adriatic Balkans were included in the phylogenetic delineation, (Figure 3d).

**Figure 3.**
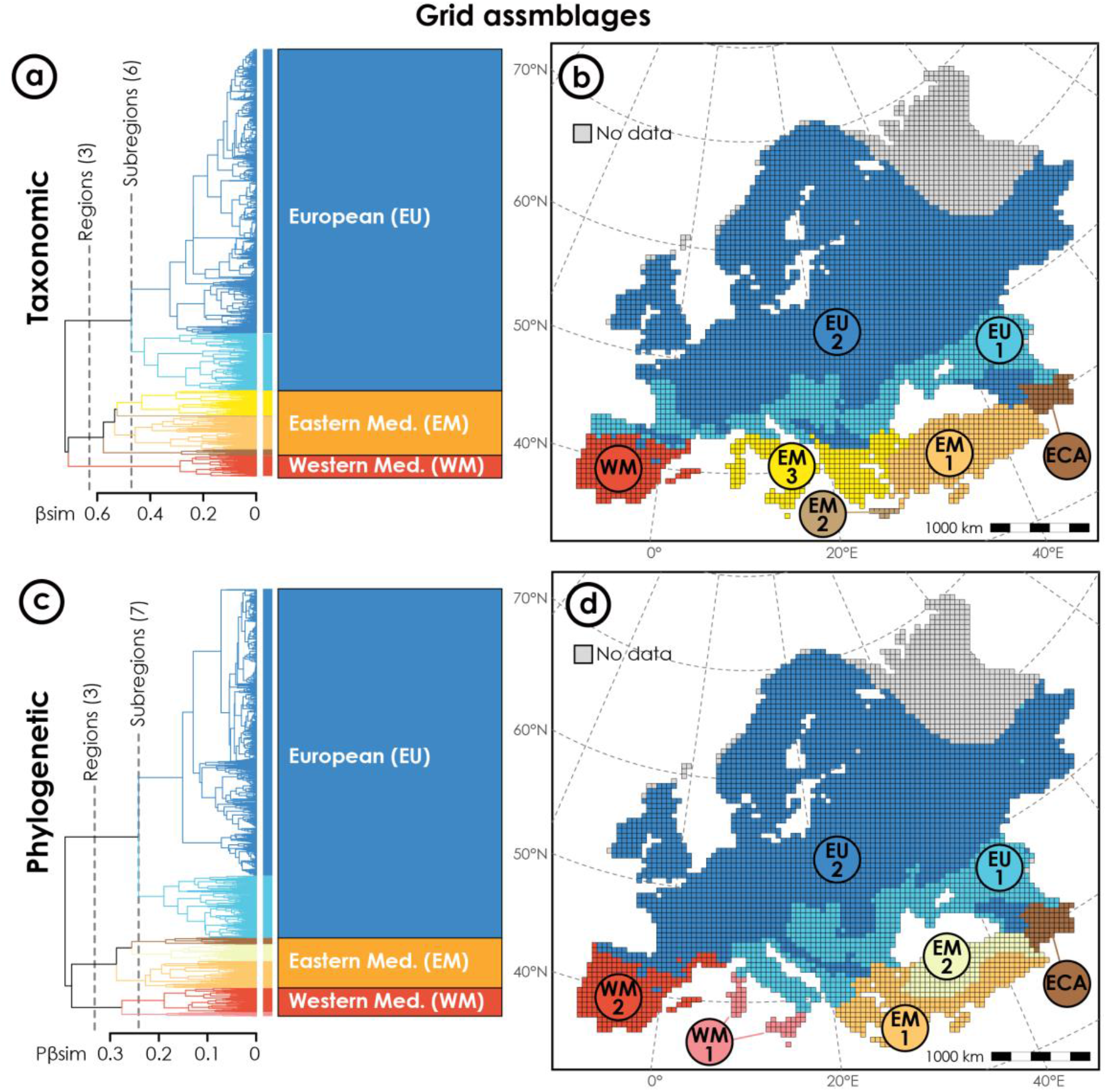
Regionalization of the western Palearctic ant fauna based on grid assemblages. Dendrograms (a, d) and maps (b, e) resulting from UPGMA hierarchical clustering based on βsim (a-c) and Pβsim (d-f) matrices. Colours used to characterize particular regions in dendrograms and maps are identical. Values that are not available were indicated by the grey colour on the map.

#### Biogeographic boundaries

Most of the boundaries of biogeographic regions and subregions of western Palearctic ants were along the mountain chains or other geographic barriers like seas (Figure 4). The Pyrenees, Apennines, Dinaric Alps, Balkan mountains, Black Sea, and the Caucasus mountains separated the southwestern (i.e., Mediterranean regions) and northwestern (i.e., European region) Palearctic realm of ants, from west to east. Mediterranean regions were mainly divided by the Mediterranean, Tyrrhenian and Ionian Seas while the Mediterranean and Aegean Seas, Taurus and Lesser Caucasus mountains were the major boundaries of subregions. The boundary of subregions in the European region was also located along mountain ranges including the Alps, Carpathians and Great Caucasus. The phylogenetic approach detected more boundaries in the Eastern Mediterranean region compared to taxonomic delineation (Figures 4b and c). Biogeographic boundaries identified based on grid assemblages were more consistent across dissimilarity metrics and better matched geographic barriers compared to the boundaries estimated using regional lists. And boundaries delineated using both geographic units were well-matched when the regional polygons are delimited along mountains, for example, on the Balkans, Taurus and Caucasus mountains (Figures 4b and c).

**Figure 4.**
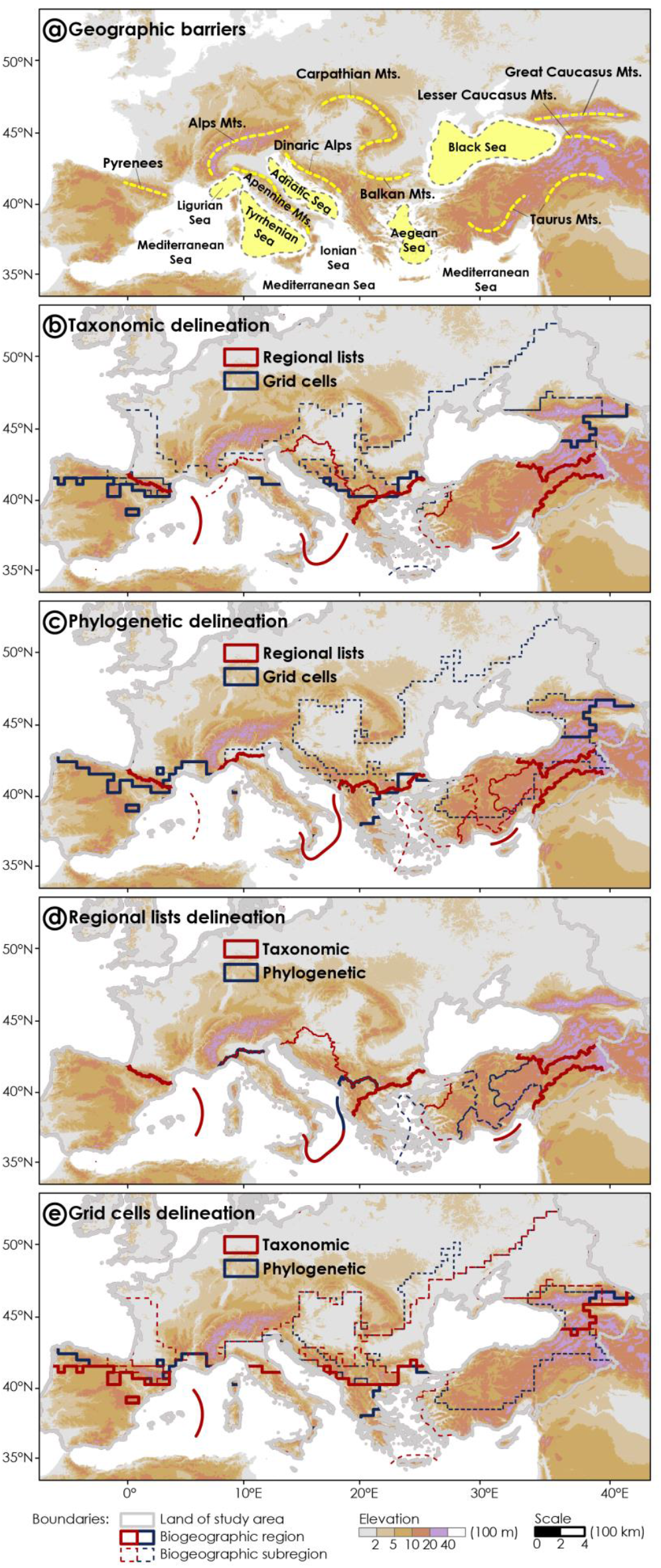
Comparison of biogeographic structure transitions. The major geographic barriers (a) are indicated by the yellow dotted lines. The boundaries of biogeographic regions/subregions based on regional lists and grid cells are overlapped in taxonomic (b) and phylogenetic (c) delineations while the boundaries of taxonomic and phylogenetic delineations are also compared at regional list- (d) and grid cell-level (e). The biogeographic boundaries over seas are also indicated.

## DISCUSSION

In this study, we proposed a new and comprehensive regionalization system for the western Palearctic ants. To the best of our knowledge, this novel system represents the first comprehensive delineation for Palearctic insects at a fine geographic scale informed by a large-scale phylogeny. The regionalization of ants proposed in this study largely follows the typical biogeographic structure of western Palearctic fauna, but at the same time reveals differences across geographic units and dissimilarity metrics.

### Biogeographic regionalization of ants in the western Palearctic realm

Overall, our results show that the western Palearctic ant fauna has a clear biogeographic structure: it can be separated into southwestern (i.e., Mediterranean regions) and northwestern parts (i.e., European region), with the southwestern part having stronger regionalization (i.e., higher turnover and more biogeographic regions) compared to the northwest (Figures 1-3). This biogeographic divergence of the western Palearctic realm is consistent with the first zoogeographic regionalization proposed by Wallace (1876) and other systems proposed afterwards based on vertebrates (e.g., Holt et al., 2013; Ficetola et al., 2018) and insects (e.g., Heiser et al., 2014; Vitali & Schmitt, 2017).

This biogeographic divergence of ants reveals the strong impact of historical climate changes on species distributions in the western Palearctic. The comparison of fossil and modern ant assemblages has shown that many lineages went extinct in the western Palearctic realm during the cooling period following the Miocene (Guénard, Perrichot & Economo, 2015). As more diverse and unique ant assemblages currently exist in Mediterranean regions, perhaps due to higher climate stability (Schmitt, 2007), this suggests that the historical signal of western Palearctic ant fauna may persist in these areas (Figure 1). In contrast, the homogenous structure of ants in the northwestern Palearctic realm may indicate that the fauna in the north was assembled by poleward dispersals of species from their southern glacial refugia after the Last Glacial Maximum (Baroni Urbani & Collingwood, 1977; Pusch et al., 2006; Leppänen et al., 2013), a pattern which is also found in many other taxa (Dennis et al., 1991; Fattorini & Ulrich, 2012; Vitali & Schmitt, 2017). The Pyrenees, Balkans and Caucasus are likely to be important postglacial recolonization centres for ants due to the strong biogeographic affinity between these regions and the northwestern Palearctic realm (Figures 3 and S2.7).

The strong biogeographic regionalization in Mediterranean regions also suggests the importance of dispersal limitation in shaping ant biogeographic structure. Compared to the northern part of the western Palearctic realm, Mediterranean regions have more islands and varied topography, which may be responsible for restricting ant distributions (Figure 4). Especially for the East Mediterranean region, its remarkable uniqueness in species composition and biogeographic isolation could also be the consequence of complex topography and geological history (Vitali & Schmitt, 2017; Ficetola et al., 2018; Ahmadi et al., 2021; Kiran & Karaman, 2021). Thus, the dispersal limitation may explain why geographic barriers (e.g., mountains and seas) represent major boundaries separating biogeographic regions and subregions of the western Palearctic ants (Ficetola, Mazel & Thuiller, 2017).

Moreover, the strong biogeographic affinities between some isolated regions also reveal the legacy of historical land connections. The high similarity in the fauna of the Apennine Peninsula and Adriatic Balkans, the Corsica and Sardinia islands, and the Sicilian and Maltese islands (Figures 2 and 3) can be explained by the biotic exchanges through land bridges during the period of partial disappearance of the Mediterranean Sea in late Miocene (i.e., Messinian salinity crisis) and Quaternary glacial periods (Randi, 2007; Schlick-Steiner et al., 2007; Schmitt, 2007; Dapporto et al., 2014; Schmitt et al., 2021). The biogeographic affinity of ant assemblages between the Tyrrhenian islands and the Iberian Peninsula suggests the evolutionary history shared by their fauna (Figures 2c-d and 3c-d). This may be due to the historical expansion of old lineages through land bridges (Senczuk et al., 2017; Dapporto et al., 2019; Schmitt et al., 2021), or to the legacy of more ancient geographic connections. Corsica and Sardinia islands were parts of the Tyrrhenian microplate which was separated from the Iberian Peninsula, and thus some lineages may persist with the same origins as the Iberian fauna (Ketmaier, Caccone & Silva-Opps, 2013; Schmitt et al., 2021).

### Effects of geographic units and dissimilarity metrics on regionalization

#### Regional lists versus grid cells

Both regional lists and grid assemblages result in a very similar biogeographic structure, although several species are not included in grid-level analysis (Figures 2, 3 and S2.5). The regionalization based on grid assemblages shows more detail regarding biogeographic transition and seemingly detects stronger effects of geographic barriers in comparison to that of the regional lists. The grid-level delineations reveal not only the close relationship between the Pyrenees, Balkans, Caucasus and the northwestern Palearctic ant fauna, but also suggest differences between the southern subregion and the European region (EU in Figure 3). The southern subregion (i.e., EU1 in Figure 3) includes several glacial refugia (Hewitt, 1999; Schmitt, 2007) and may harbor species extinct in the north during the Last Glacial Maximum and unable to disperse since across physical barriers like the Alps and the Carpathian mountains (Figures 4 and S2.8).

The accuracy of regional polygons would influence the regionalization results. Due to their relatively coarse scale, regional polygons likely do not reflect species’ range boundaries as accurately as grid cells (Figure 4). Some regional polygons based on administrative divisions may extend across multiple biogeographic regions and thus regional lists may include distinct fauna. For example, the regional list of southeastern Anatolia has a relatively large extent that may include some species from the Arabian Peninsula, making its fauna very different from the rest of the Eastern Mediterranean region. Notably, delineations of regional lists highlight the biogeographic uniqueness of Sicily, Maltese and Cyprus islands (Figure 2). The SDMs we used to make range predictions do not correct for dispersal limitations, so the long-term isolation and complex geologic history of these islands (Poulakakis et al., 2013; Schmitt et al., 2021) and other regions with similar histories were not considered for the grid assemblage regionalization.

#### Taxonomic versus phylogenetic metrics

The spatial patterns of taxonomic and phylogenetic turnover were highly congruent (Figure 1 and Table S2.3). This suggests that the spatial turnover of ant phylogeny may be caused by species replacement in the western Palearctic realm, while the null model analysis suggests many coastal and insular Mediterranean regions present strong phylogenetic turnover which is independent of species richness and compositions. The phylogenetic regionalization further reveals the affinity between the Tyrrhenian islands and the Iberian Peninsula and the divergence between northern and southern Anatolia (Figures 2c-d and 3c-d); the latter is perhaps even under-estimated due to the limitation of phylogenetic data (Figure S2.4). Thus, the phylogenetic approach provides evolutionary support and some new perspectives into the biogeographic history of the West Palearctic realm.

### Tackling the Wallacean and Darwinian shortfalls for diverse insect groups

Answering some of the oldest and most important questions in biology relies on basic information about species distributions and their phylogenetic relationships, which represent important shortfalls for most insect groups. These knowledge gaps have long prevented biogeographic classifications based on taxa other than plants and vertebrates. In particular, classifications made based on vertebrates are particularly problematic, as current data may hide profound assemblage changes that occurred following ancient human settlement and resulting environmental modifications (Faurby & Svenning, 2015; Santos et al., 2020). As a result, datasets that expand the taxonomic breadth of biogeographic classifications are needed to confirm previous results if any broad patterns across taxa are to be confirmed. Although rare for insects, this study was able to produce a comprehensive regionalization of the western Palearctic ants due to the development of accurate and exhaustive distributional information, as well as a large-scale phylogeny.

This study demonstrates that a combination of expert opinion and modeling can help tackle the Wallacean Shortfall for insects. The expert-validation step ensures quality control of the distributional data, while models like SDMs represent powerful tools for estimating detailed delimitations of species distributions. However, we also recognize some limitations in this approach. For example, the SDM predictions we used here represent the responses of ant species to climatic variables and do not explicitly consider species’ dispersal abilities, biotic interactions, and other local effects. Although techniques exist to consider dispersal limitations (Monsimet et al. 2020) and biotic interactions (Kass et al. 2020, Wisz et al. 2013) in SDMs, this is currently difficult for ants because the necessary data does not exist for most ant species. Thus, the range estimates from the regional lists and the SDM predictions can suggest different affinities and spatial extents in biogeographic structure (Figures 2, 3 and S2.7). Such important ecological information could improve the modeling of species distributions and the resulting delineation of biogeographic structure once available for more ant species (e.g., Jaeschke et al., 2013).

The Darwinian Shortfall for insects may be particularly important for many regions around the globe. Some of the diverse insect groups for which large-scale phylogenies have been developed could be ideal models for understanding the macroevolution and macroecology of insects (e.g., ants: Economo et al., 2018, butterflies: Earl et al., 2021). The estimates of phylogeny could be useful for solving limitations due to limited molecular data available in most insect groups, especially for metrics like phylogenetic turnover which is based on the branches length, and are not sensitive to the uncertainty in phylogenetic topology (Economo et al., 2018; Jetz & Pyron, 2018). However, more taxonomic revisions and molecular sequencing are needed to complete comprehensive and robust phylogenetic constructions for insect species.

## Conclusion

Our study shows that expert validation and modeling of species distributions combined with a large-scale phylogeny can help us develop a comprehensive regionalization for a diverse insect group, and thus directly tackle Wallacean and Darwinian shortfalls for insects. The new bioregionalization of western Palearctic ants we present here supports a biogeographic divergence between the more homogenous northwest European region and the more regionalized southwest Mediterranean region. These biogeographic structures reveal potential effects of Quaternary climate changes, and even deeper geological processes like plate tectonics, on ant distributions. The western Palearctic ants represent an ideal system to investigate how the insect distributions have responded to historical processes. Moreover, the remarkable uniqueness of coastal and insular Mediterranean areas highlights their historical roles as glacial refugia and potential significance for the future conservation of ant diversity. The estimation of regionalization systems for other insect groups would provide the possibility for comparisons with our delineations, which would contribute to essential knowledge of insect biogeography and would provide important information about their uniqueness to guide future conservation efforts.

## Acknowledgements

RW is supported by an Early Career Scheme Grant from the Research Grants Council (ECS-27106417) of the Hong Kong Government. JMK is supported by the Japan Society for the Promotion of Science (JSPS) Postdoctoral Fellowships for Foreign Researchers Program; AR is supported by the grant NRFU (Ukraine) No. 2020/02/0369. This work can not be done without the close and extensive collaborations across borders, we want to thank the peace we had in Eurasia and hope we will still have it in the future!

## DATA AVAILABILITY STATEMENT

The regional lists, binary range maps and metadata of species distribution modelings used for this study will be available in Dryad Digital Repository once the study has been accepted.

## SUPPLEMENTARY

**Appendix 1**.

**Table S1.1**. Information of species included in the study.

**Appendix 2**.

**Table S2.2**. Evaluation of clustering algorithms.

**Table S2.3**. Correlation of taxonomic and phylogenetic pairwise dissimilarity matrix and spatial turnover.

**Figure S2.1**. Diagram of the workflow of this study.

**Figure S2.2**. Map of geographical units used in the European Ant Distribution (EUAD) database.

**Figure S2.3**. Contribution of experts to the new database.

**Figure S2.4**. Spatial pattern of species without species distribution modellings or phylogenetic data.

**Figure S2.5**. Evaluation of UPGMA hierarchical clustering of ant regional lists.

**Figure S2.6**. Evaluation of UPGMA hierarchical clustering of ant grid assemblages.

**Figure S2.7**. Different subregions identified within the European region of ants.

---

**Table S2.3.**
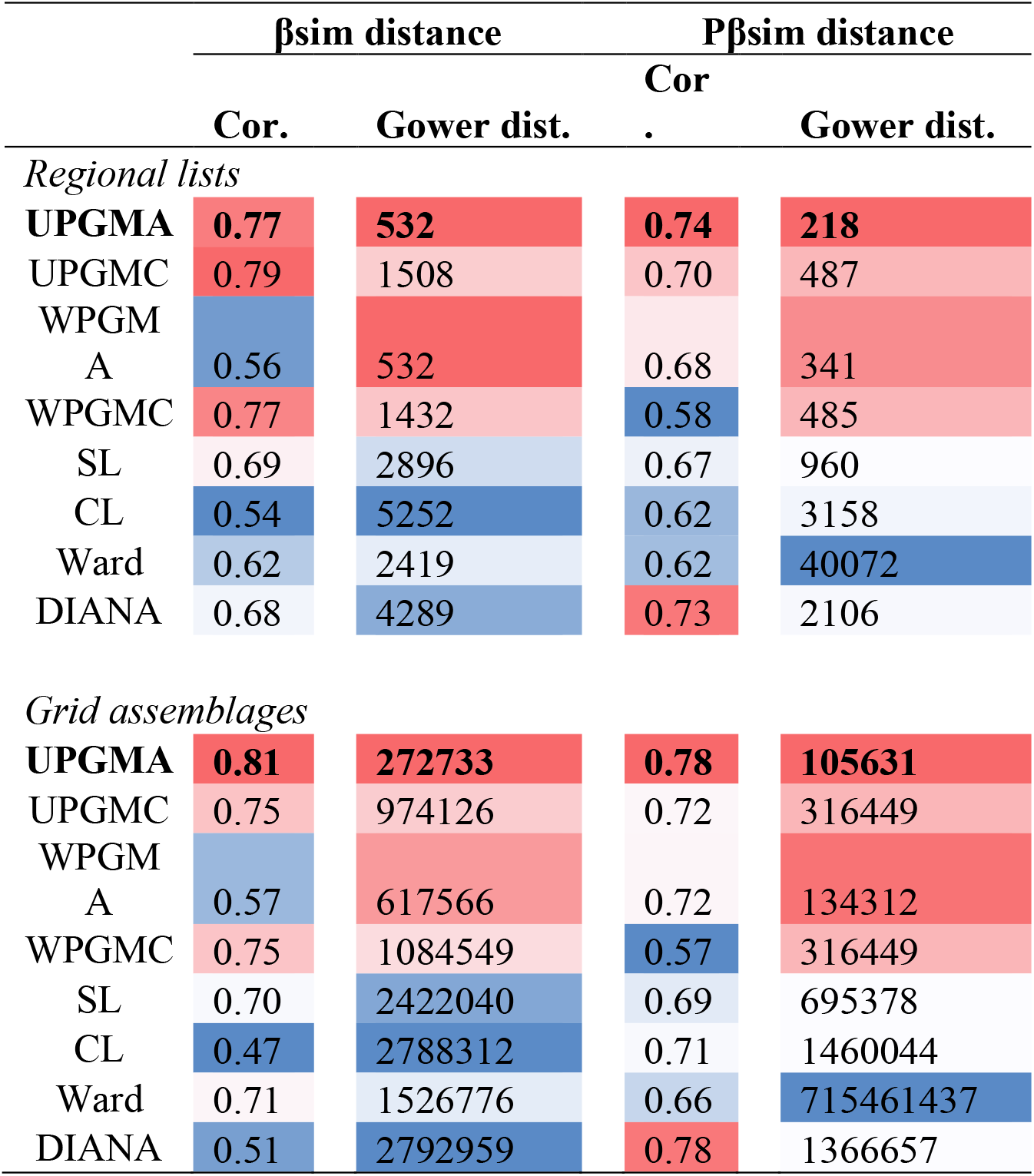
Evaluation of clustering algorithms. Algorithm with higher cophenetic Pearson correlation (Cor.) and lower Gower distance (Gower dist.) is considered to have a better performance. Red and blue colours indicate better and worse performance in each criterion, respectively. Abbreviations: UPGMA, unweighted pair-group method using arithmetic averages; 2) UPGMC, unweighted pair-group method using centroids; 3) WARD, Ward’s method; 4) SL, single lineage; 5) CL, complete lineage; 6) WPGMA, weighted pair-group method using arithmetic averages; 7) WPGMC, weighted pair-group method using centroids; 8) DIANA, divisive hierarchical clustering.

**Table S2.4.** Correlation of taxonomic and phylogenetic pairwise dissimilarity matrix and spatial turnover. Pearson’s correlation assessed by Mantel test is shown above the diagonal and spatially corrected Pearson’s correlation calculated by modified t test is shown below the diagonal. All correlations are significant at *P* < 0.001 level.

| | | βsim | SES.βsim | Pβsim | SES.Pβsim |
| --- | --- | --- | --- | --- | --- |
| Regional<br>assemblage<br>s (N = 207) | βsim | — | 0.87 | 0.94 | 0.85 |
| | SES.βsim | 0.93 | — | — | 0.95 |
| | Pβsim | 0.96 | — | — | 0.82 |
| | SES.Pβsim | 0.90 | 0.94*** | 0.88 | — |
| Grid<br>assemblage<br>s (N =<br>4403) | βsim | — | 0.88 | 0.95 | 0.84 |
| | SES.βsim | 0.92 | — | — | 0.95 |
| | Pβsim | 0.96 | — | — | 0.82 |
| | SES.Pβsim | 0.91 | 0.97 | 0.85 | — |

**Figure S2.1.**
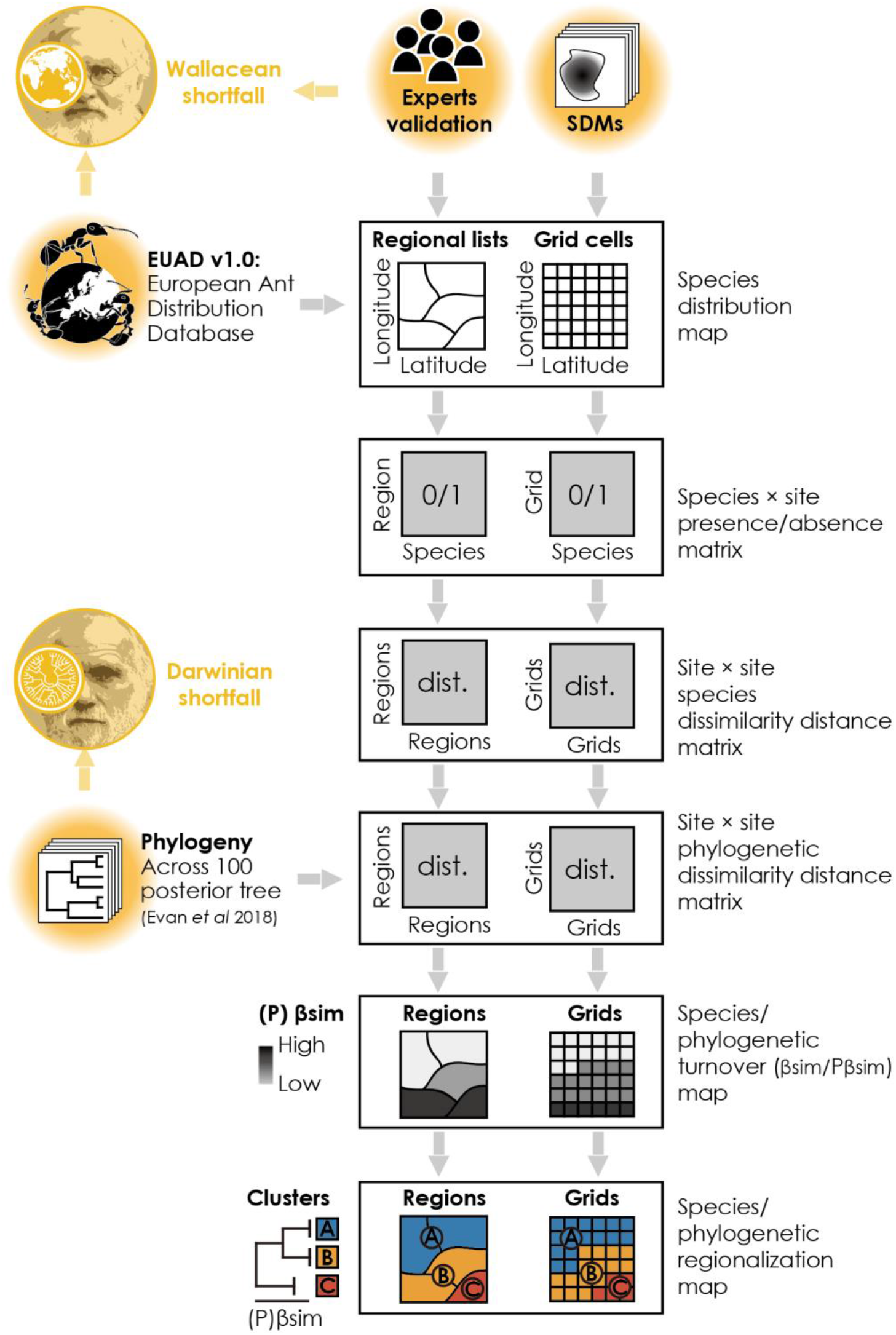
Conceptual diagram of the workflow of this study.

**Figure S2.2.**
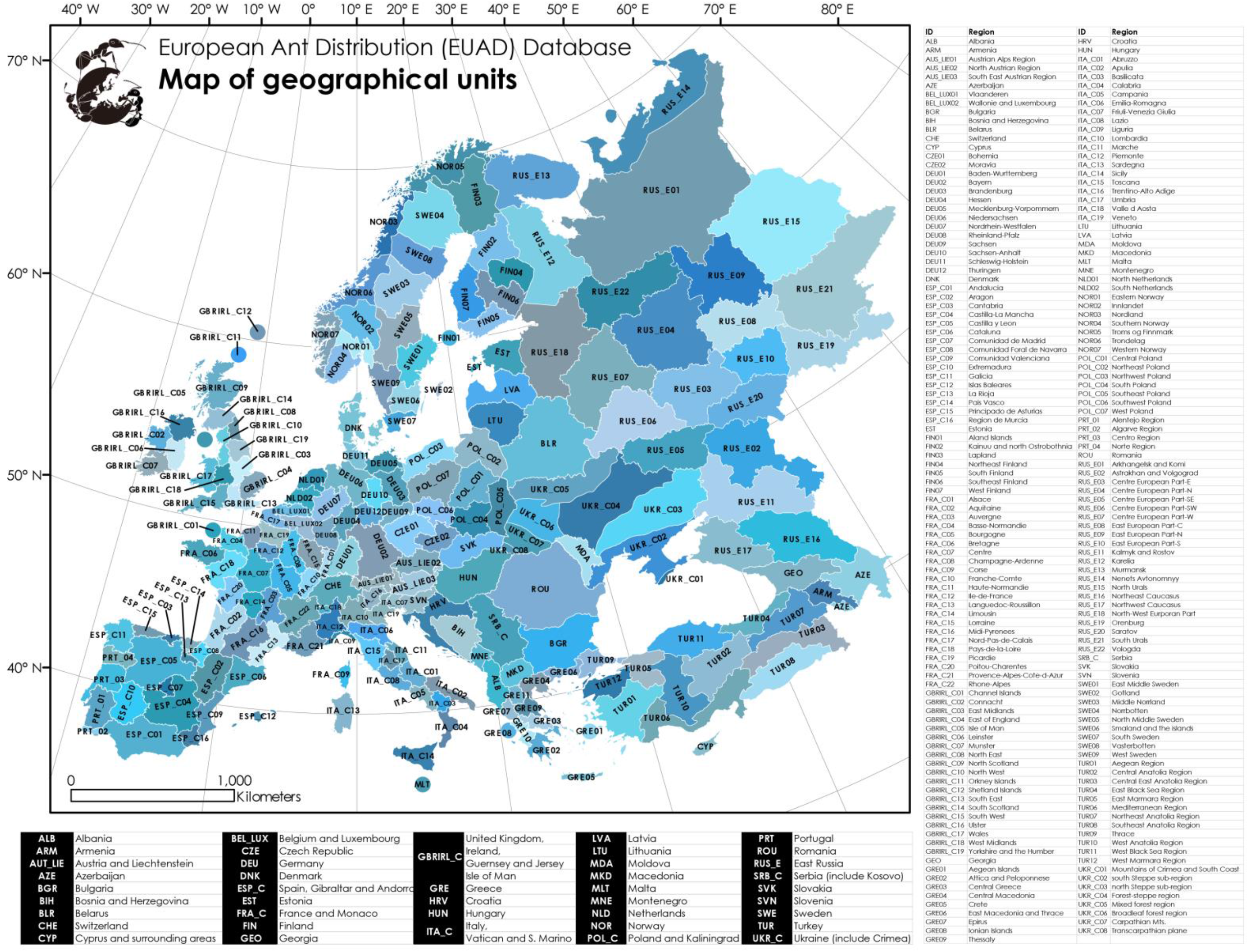
Map of geographical units used in the European Ant Distribution (EUAD) database. Abbreviations indicate the administrative or geographic regions delimited in the EUAD database.

**Figure S2.3.**
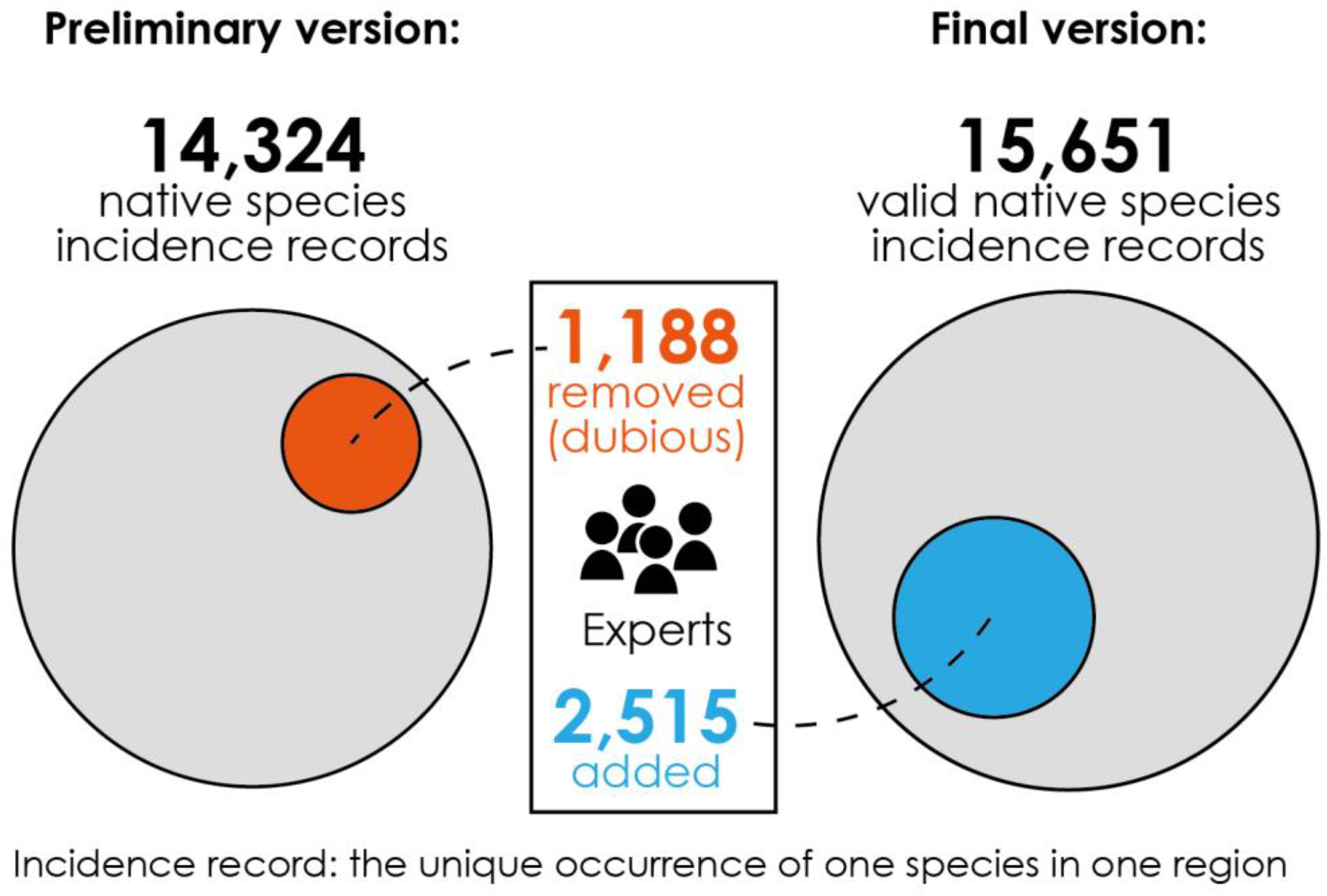
Contribution of experts to the new database. Dubious and new records are removed and added based on experts’ opinions, respectively.

**Figure S2.4.**
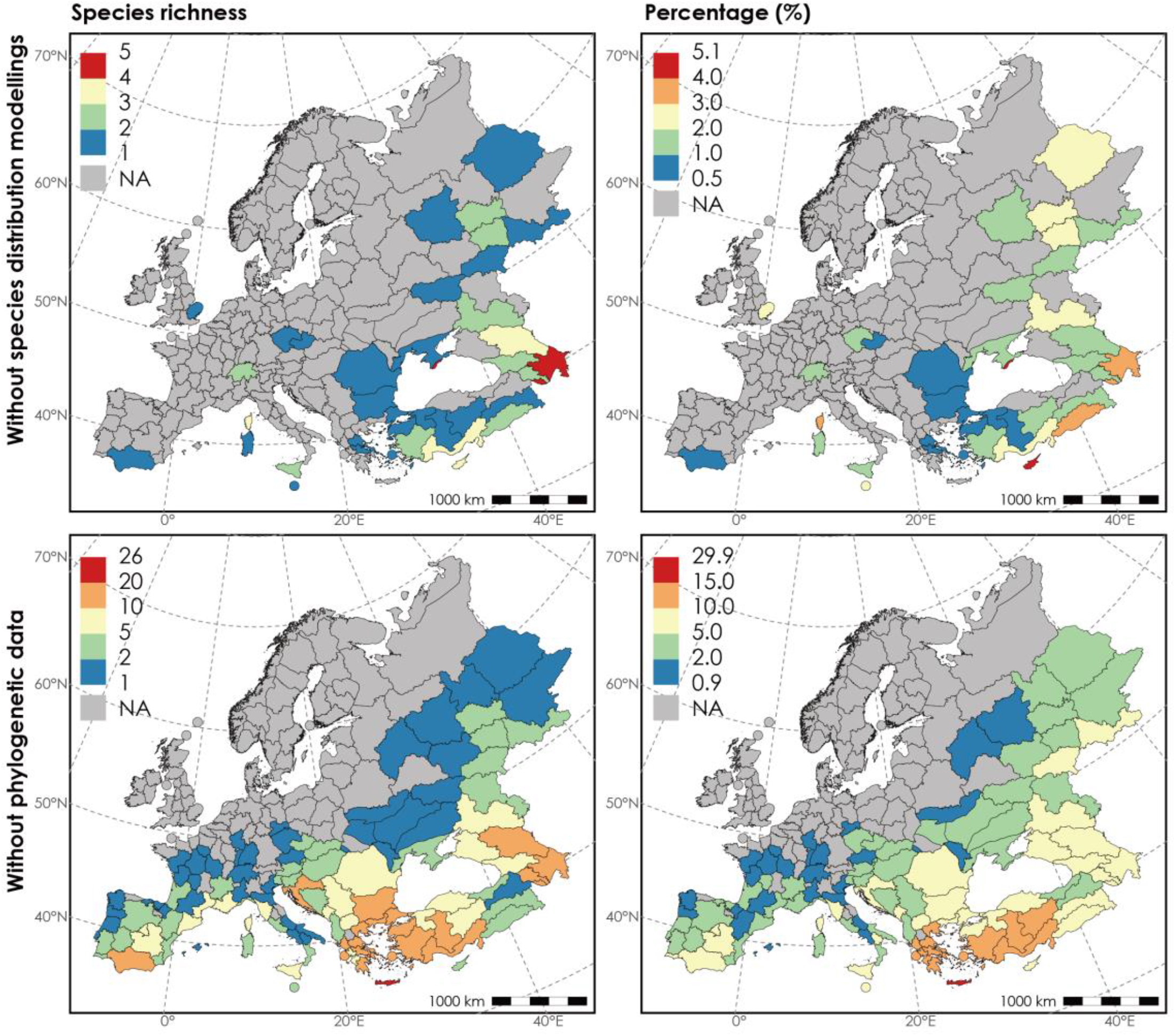
Spatial pattern of species without species distribution modellings or phylogenetic data. Both richness and the percentage of missing species are calculated for each region.

**Figure S2.5.**
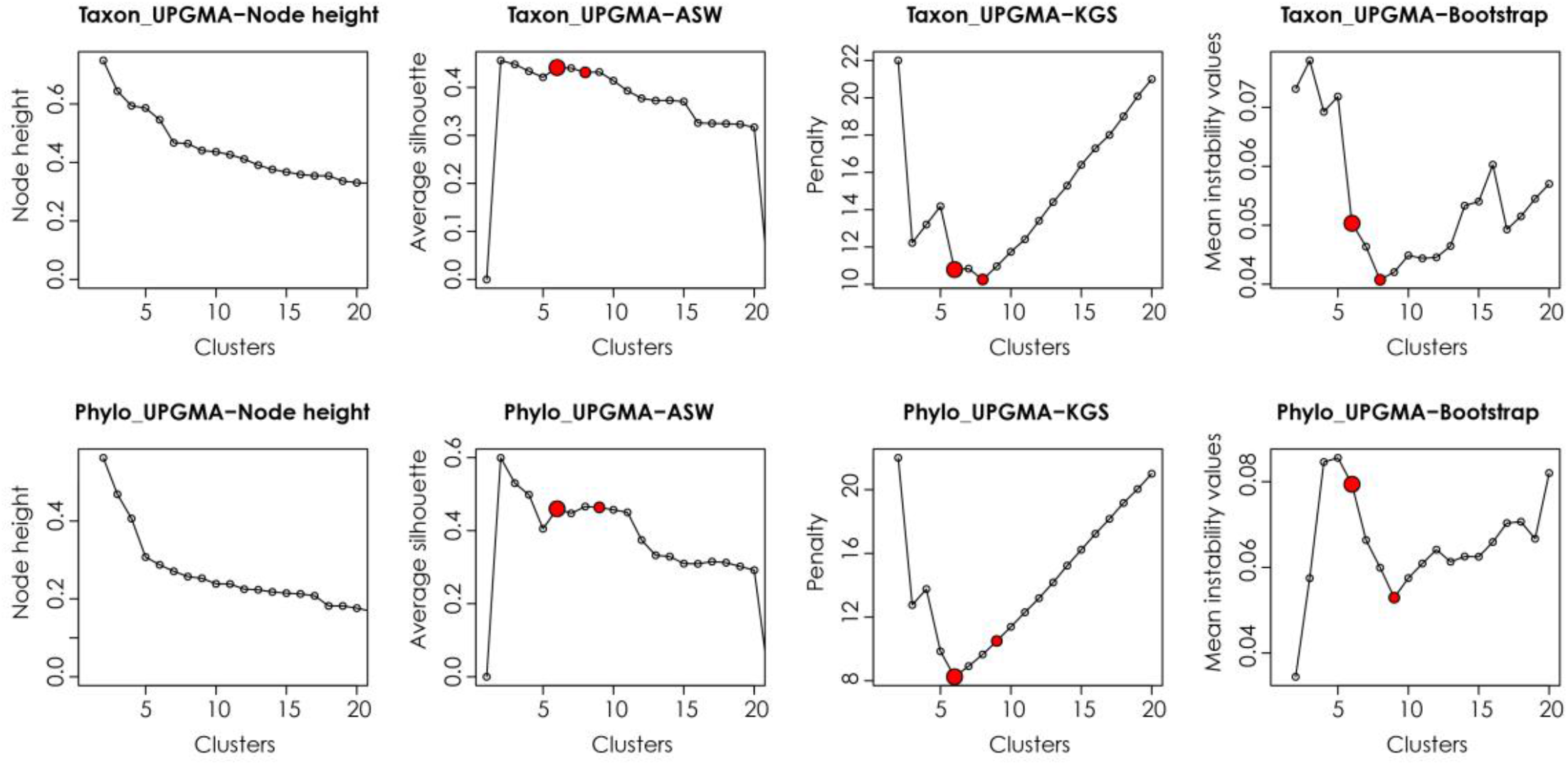
Evaluation of UPGMA hierarchical clustering of ant regional lists. The node height, average silhouette (ASW), values of Kelly-Gardner-Sutcliffe penalty (KGS) and mean instability values of Bootstrap test are calculated based on taxonomic (upper row) and phylogenetic (lower row) dissimilarity. The red dots indicate the number of clusters chosen for biogeographic regions (large dots) and subregions (small dots).

**Figure S2.6.**
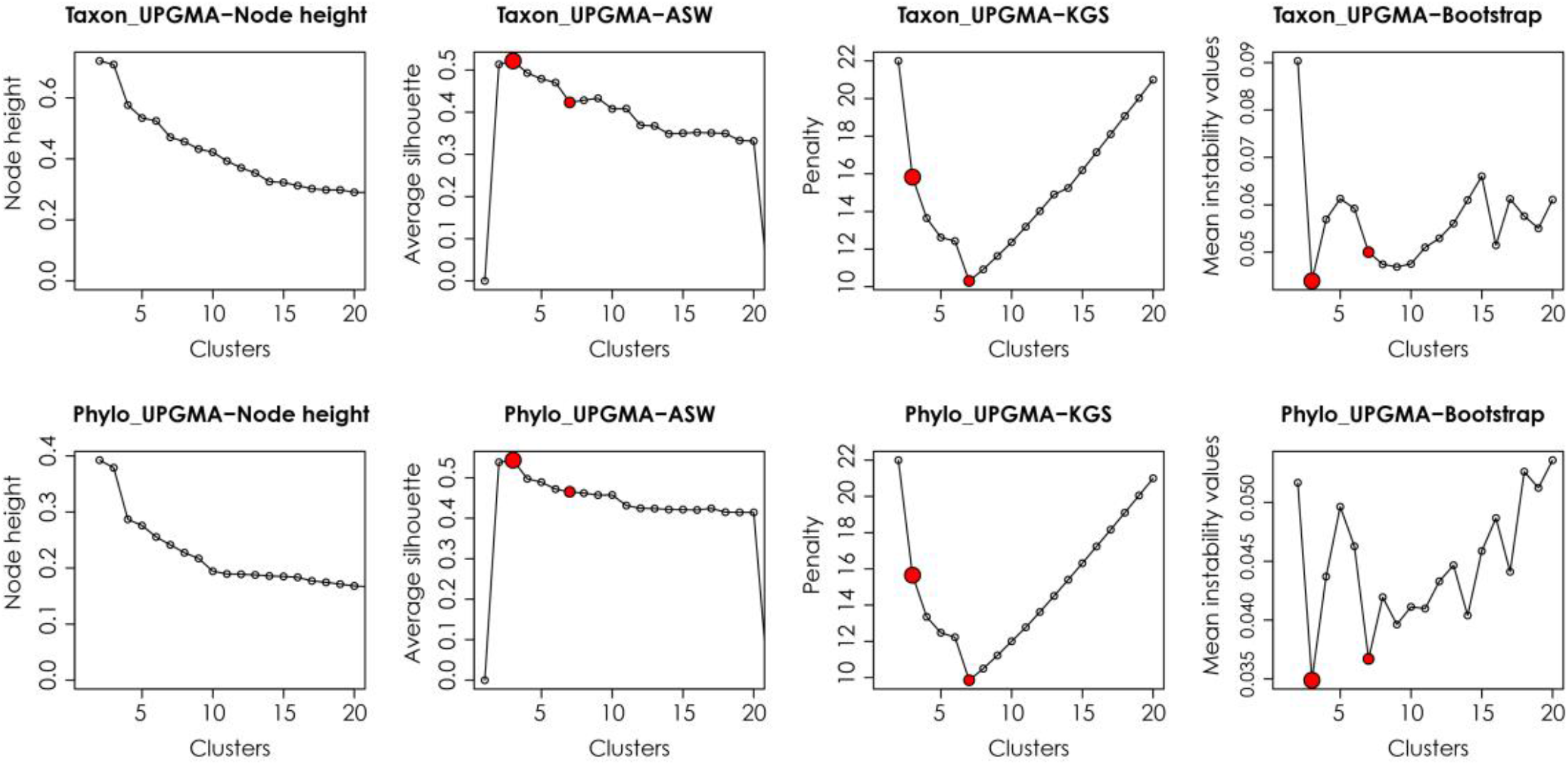
Evaluation of UPGMA hierarchical clustering of ant grid assemblages. The node height, average silhouette (ASW), values of Kelly-Gardner-Sutcliffe penalty (KGS) and mean instability values of Bootstrap test are calculated based on taxonomic (upper row) and phylogenetic (lower row) dissimilarity. The red dots indicate the number of clusters chosen for biogeographic regions (large dots) and subregions (small dots).

**Figure S2.7.**
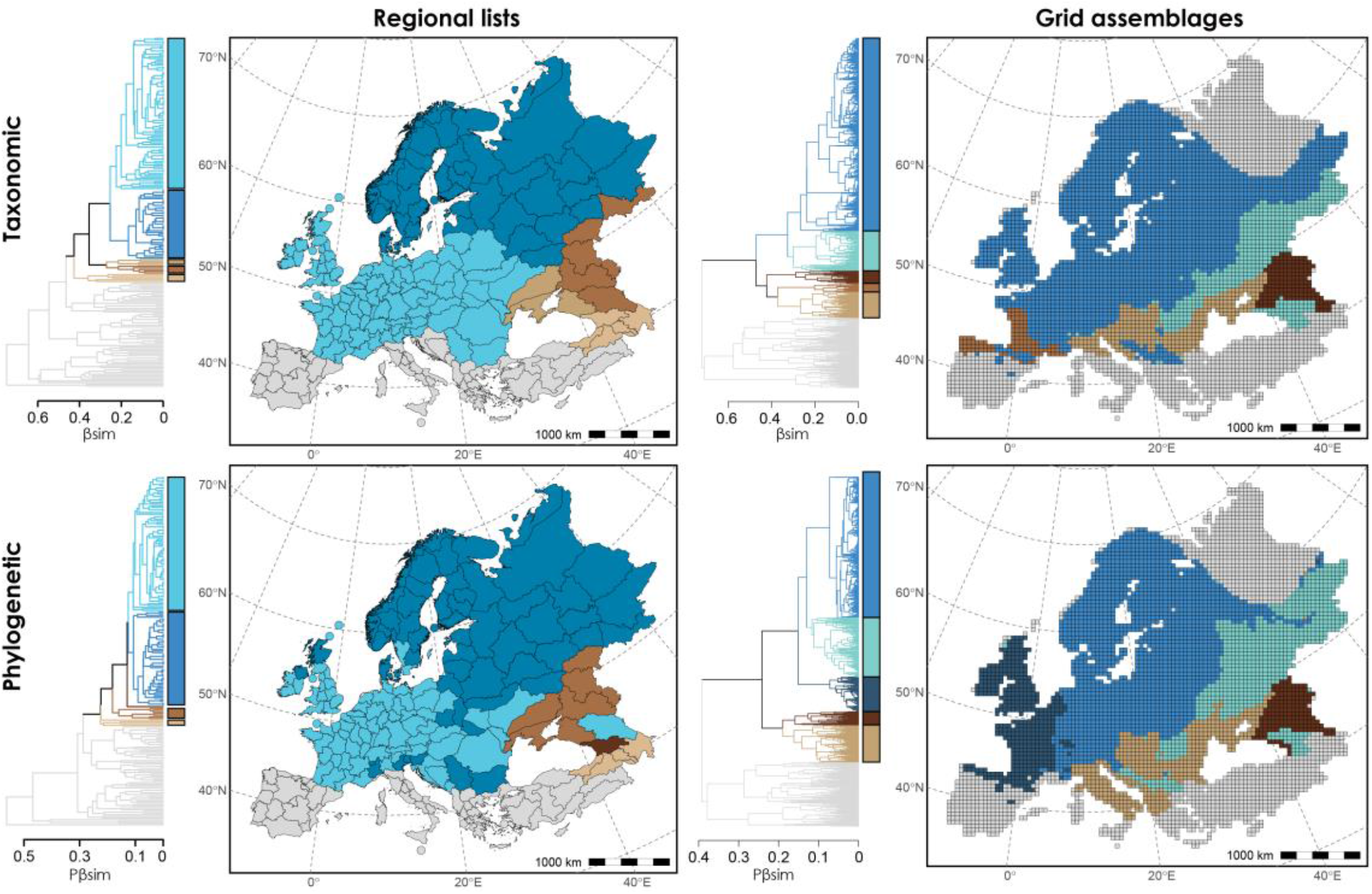
Different subregions identified within the European region (EU) of ants. Subregions are delineated based on different geographic units (i.e., regional lists and grid cells) and approaches (i.e., taxonomic and phylogenetic).

